# Oscillatory brain activity links experience to expectancy during associative learning

**DOI:** 10.1101/2021.01.04.425296

**Authors:** Kierstin Riels, Rafaela Campagnoli, Nina Thigpen, Andreas Keil

**Affiliations:** University of Florida; Biomedical Institute, Universidade Federal Fluminense, Niterói, RJ, Brazil; Department of Neurobiology, Institute of Biology, Universidade Federal Fluminense, Niterói, RJ, Brazil

**Author notes:** Corresponding Author Kierstin Riels.

## Abstract

Associating a novel situation with a specific outcome involves a cascade of cognitive processes, including selecting relevant stimuli, forming predictions regarding expected outcomes, and updating memorized predictions based on experience. The present manuscript uses computational modeling and machine learning to test the hypothesis that alpha-band (8-12 Hz) neural oscillations are involved in the updating of expectations based on experience. Participants learned that a visual cue predicted an aversive loud noise with a probability of 50 percent. The Rescorla-Wagner model of associative learning explained trial-wise changes in self-reported noise expectancy as well as alpha power changes. Both experience in the past trial and self-reported expectancy for the subsequent trial were accurately decoded based on the topographical distribution of alpha power. Decodable information during initial association formation and contingency report recurred when viewing the conditioned cue. Findings support the idea that alpha oscillations have multiple, simultaneous, and unique roles in association formation.

## Introduction

A principal function of the human brain is to build internal representations of the world, optimizing the prediction of behaviorally relevant outcomes (e.g., Hohwy, 2017). For example, the formation of food preferences involves the association of pleasant, or unpleasant, outcomes with eating a given food. Forming these associations is characterized by uncertainty, resulting in the updating and changing of associative memories (Tzovara et al., 2018). The present study combines methods from computational and cognitive neuroscience (Kriegeskorte & Douglas, 2018) to define the neural processes linking new experiences to changing expectancies. Participants were exposed to Pavlovian (Classical) conditioning, a type of learning in which an initially neutral cue (the conditioned stimulus, CS) comes to elicit an aversive or appetitive response, through association with an intrinsically relevant stimulus (the unconditioned stimulus, US; Pavlov, 1897; Lattal, 2013). The principles of Pavlovian conditioning have been extensively used to explain and predict the formation of associative memories, assessed using behavioral, cognitive, and physiological responses (Maren & Quirk, 2004). The fundamental nature and wide applicability of Pavlovian conditioning have prompted the development of computational models (Siegel and Allen, 1996). One of the most widely used models is the Rescorla-Wagner (RW) learning model, which emphasizes learning from surprising, unpredicted US occurrences.

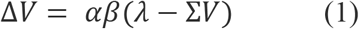

In this model, subjective “surprise” is quantified in a given experimental trial by the difference between the outcome in this trial (*λ*) and the outcome expected based on previous trials (Σ*V*) referred to as the prediction error (*λ* − Σ*V*). The learning rate parameter (*α*) represents the speed of updating the learned associative strength (*V*) based on the prediction error: As surprise increases, changes in associative strength (Δ*V*) are larger (Rescorla and Wagner, 1972; see eq. 1). Thus, in a basic implementation with one conditioned stimulus, the current associative strength is the sum of the associative strength Σ*V* in the prior trial, and Δ*V*. Conceptually, associative strength has been linked to expectancy of the US, i.e. heightened associative strength is reflected in greater expectancy of the US when presented with the CS. Thus, in the case of an aversive US, these changes can be indexed by self-report, but also by an array of autonomic and reflex measures of defensive activation (Hamm & Weike, 2005). Conditioned stimuli may also differ in terms of saliency, which are modeled by the *β* parameter. In the original version of RW, this parameter is inextricable as it is multiplicatively associated with the learning rate (*α*). Thus, these parameters are typically considered together in the original version of the RW model.

Associative strength is often conceptualized as the extent to which conditioned and unconditioned cues are linked, measured by the capacity of the conditioned stimulus to evoked the conditioned response. The extent of trial-by-trial changes in associative strength is limited by the learning rate, assumed to be constant at a given time and in a given person (Rescorla & Wagner, 1972). Additional models have been widely used, notably variants of the Pearce-Hall model (Pearce & Hall, 1980) and the Mackintosh model (Mackintosh, 1975), both of which can be considered extensions of the RW model that allow for the learning rate to be dynamically updated as learning progresses (Mitchell & Pelley, 2010), often using an additional variable, related to the acquired saliency, attention or “associability” of the stimulus, linked to its past surprise (Li et al., 2011). More recent models have built on these developments and also account for changing CS and US saliency within each trial, allowing researchers to fit this parameter separately from the learning rate (Esber & Haselgrove, 2011; Kruschke, 2001).

The two main variables of the RW model, associative strength (*V*) and prediction error (*λ* − Σ*V*), have been related to a range of empirical indices traditionally used to measure association formation, such as various EEG, fMRI-BOLD, and behavioral changes throughout an experiment (Williams et al., 2017; Roesch et al., 2012). Fitting associative strength and prediction error parameters to neural time series while estimating parameters of the RW or related models has the advantages of productively using the trial-by-trial variability in the brain data, combined with conceptually meaningful data reduction, and the ability to test specific quantitatively stated hypotheses. Accordingly, this approach has been used to quantify learning in a wide range of tasks, in human participants and animal model studies, including reinforcement learning (Costa et al., 2016), social decision making (Zhang et al., 2020), and category learning (Rodriguez et al., 2006). The present study aims to use this technique to characterize the role of alpha-band oscillations during experience-based updating of expectancy.

EEG data represent an index of cortical activity with high temporal fidelity. Brain oscillations extracted from EEG are traditionally categorized into frequency ranges that have been linked to broad behavioral states. Changes in spectral power in the alpha band (8-12 Hz) are reliably associated with specific cognitive processes (Klimesch, 1999). Internally focused cognitive processing, such as imagery (Bartsch et al., 2015), boredom (Raffaelli, Mills, & Christoff, 2018), and anticipation (Bonnefond & Jensen, 2012), have been linked to alpha power increases. In contrast, cortical engagement with relevant and/or salient sensory input is associated with alpha power decreases (Berger, 1929; Klimesch, 1999). Together, this prompts the hypothesis that updating learned representations, which involves anticipation, stimulus selection, and memory re-formation involves dynamic changes in alpha-band neural activity. Testing this hypothesis is facilitated by the fact that alpha-band oscillations are high in amplitude and possess favorable signal-to-noise ratio, allowing researchers to reliably quantify them at the single-trial level, and over each time point in a trial (Başar & Güntekin, 2012).

Stimulus saliency and changing expectancy are two keys factors linking associative strength in the RW model to neural oscillations in the alpha band. When a neutral stimulus is reliably paired with a relevant outcome, behavioral and physiological responses to this conditioned stimulus change in predictable ways, optimizing the readiness of the organism for the predicted outcome (Clark, 2004). Importantly, these responses include heightened sensory responses to conditioned cues, consistent with increasing sensory saliency and heightened attention (e.g., Miscovic & Keil, 2012). Expectancy is likewise increased as the associative strength between a conditioned stimulus and unconditioned stimulus increases (e.g., Perruchet, 2015). These findings readily translate into the specific hypothesis that associative strength in the RW model is reflected in alpha power fluctuations. Because alpha power and timing are highly variable per individual (Haegens et al., 2014), a model that is built on individual learning rates and neural characterizations can create a more complete picture of associative learning than the group and trial-averaged data. The present study leverages the high temporal and spatial fidelity of multi-channel EEG recordings to model trial-wise fluctuations in time-varying alpha-band activity within the RW framework. Using model comparison and model validation, we also compare the RW model against more flexible computational frameworks for describing alpha-band fluctuations during associative learning.

In addition to the conceptually-driven RW model, the present study used purely data-driven approaches to test the hypothesis that alpha power fluctuations reflect the updating of associative strength during learning. Automatic machine classifying systems learn and detect diagnostic differences from the data, rather than from a theoretical model such as the RW framework (Vu et al., 2018). One machine learning technique, support vector machine (SVM) classification, separates observations based on a set of features. SVM classification has been widely used in neuroscience research, to characterize fMRI voxel patterns linked to neural representations of specific stimulus categories (Haxby et al., 2014), and to identify periods in which EEG scalp topographies discriminate between cognitive states (Bae & Luck, 2018). Thus, SVM classifiers allow for the identification of neural representational patterns that correspond to the cognitive and/or behavioral states challenged by the experimental paradigm (Weaverdyck et al., 2020). More specifically, temporal generalization analyses use the time information inherent in EEG signals to examine the stability and similarity of representations across time points (King & Dehaene, 2014). Implementing this approach along with the individual RW modeling approach allows us to computationally characterize the relation between alpha power fluctuations and the trial-by-trial updating of associative memories. It also allows us to test the hypothesis that the spatial distribution of alpha power across the scalp contains decodable information representing both experience and changing expectancy, and that this information is specific to different phases within a learning cycle.

### The Current Study

The present study utilizes a simple classical conditioning paradigm in which the CS, a visual grating stimulus, is occasionally (50 % contingency) paired with the US, a loud white noise burst. After each trial, participants report their expectancy of a loud noise to occur in the next trial. This design prompts high levels of uncertainty and encourages sustained updating of expectancy, throughout the experimental session (Moratti & Keil, 2009; Perruchet, 2015). The absence of additional manipulations also facilitates the interpretation of changes in the behavioral (expectancy ratings) and brain data (time-varying EEG alpha band activity) as primarily influenced by individual learning rates, prediction error, and baseline associative strength. We hypothesize that fluctuations in alpha-band power reflect associative strength and as such are modeled by the associative strength variable in the RW model (Miller et al., 1995). Given the critical role of alpha-band activity in sensory and anticipatory processing, we also hypothesize that the topographical distribution of alpha power across the scalp contains decodable information, representing experience (US vs. no-US) as well as expectancy regarding the subsequent trial. Exploratory analyses focus on identifying the time course and nature of these SVM-based representations, and how they relate to RW parameters based on behavioral (expectancy rating) data.

## Methods

### Participants

Twenty students (15 female) from the University of Florida participated in this study for course credit. Eleven participants identified as Caucasian (55%), 6 identified as Hispanic (30%), 3 identified as Asian (15%), and the mean age was 19 years (*SD* = 0.89). All participants read and signed informed consent forms before participation and received course credit as compensation. One additional participant was run but had to be excluded from analyses because of errors resulting in an incomplete recording. This study was approved by the local Institutional Review Board.

### Task

The conditioned stimulus (CS) was a Gabor patch with a 1.5 degree left tilt and a Michelson contrast of 0.63 presented for 2.5 seconds in every trial. This stimulus was presented in the center of a SONY CRT 27” Trinitron monitor set to a 60 Hz refresh rate, spanning a visual angle of 7.8 degrees, viewed from a 120 cm distance. The unconditioned stimulus (US) was a 96 dB (SPL) noise lasting 1 second, beginning 1.5 seconds after the start of the CS, and thus co-terminating with the US in noise trials. The noise was delivered through 2 computer speakers positioned behind the participant. The CS was randomly (drawn without constraints from a rectangular distribution) paired with the US in 50% of the trials and was followed by a fixation cross lasting 1 second and a blank screen lasting another second. The free randomization resulted in a median number of 60 USs (loud noises) presented to each participant (range: 51 to 66 USs). At the end of each trial, participants were prompted to use the mouse to rate, on a visual analog scale from 1 to 10, their expectancy that the noise would occur in the upcoming trial (“How likely is the loud noise to follow the pattern?”). Free randomization was used to promote uncertainty and to discourage participants from forming heuristics such as (“there is never more than three in a row”), which has been shown to affect trial-wise updating of expectancy (e.g., Perruchet et al., 2006). A fixation cross followed the response and lasted 3 seconds before the start of the next trial. There was a total of 120 trials.

### Procedure

After reading and signing the informed consent, participants sat in a sound-dampened room with dimmed lights. After instruction procedures, research assistants placed and positioned the EEG recording net over the participant’s head and made any necessary adjustments to sensor position and impedance levels before leaving the participant and initiating the task.

### Data Acquisition and Analysis

EEG data were continuously recorded through a 129-channel sensor net with a 500 Hz sampling rate. Data were collected using an Electrical Geodesic amplifier with an input impedance of 200 MΩ. Impedances were kept below 60 kΩ, appropriate for a high input impedance amplifier. The signal was collected with online elliptical filters in place at .1 Hz high-pass and 90 Hz low-pass (defined as the 3-dB points). All data were recorded to a single Cz reference and further filtering was done offline. All channels were preprocessed with 40 Hz low-pass (Butterworth, 23^rd^ order, cut-off defined as 3 dB point) and 1 Hz high-pass (Butterworth, first order, cut-off defined as 1 dB point) filters. Epochs consisted of 3.6 seconds (1800 sample points) before and 1.5 seconds (750 sample points) after CS (Gabor patch) onset. Artifacts were rejected within these epochs based on absolute value, standard deviation, and maximum of differences across time points and for every channel, as suggested by Junghöfer and colleagues (2000). On average of 76.3 out of 120 trials were kept per participant before separating data by condition.

#### Analysis of expectancy ratings

RW modeling analyses used all trial-by-trial ratings, which normalized by the maximum at the level of individual participants to minimize scaling differences and establish a common scale between model and data, with values ranging between 0 and 1. For descriptive and decoding analyses, trial-by-trial expectancy ratings were first reduced at the participant level. To this end, a tertile split was performed, and low versus high expectancy trials (lower versus upper tertiles) were identified. Thus, individual scores were categorized into low (US unlikely to occur) or high (US likely to occur) expectancy trials respectively, with the middle tertile being discarded. To allow descriptive analyses of the interaction between experience and subsequent expectancy, we also cross-tabulated the rating data based on the outcome of the previous trial (noise versus no noise) and a low versus high expectancy rating.

#### EEG post-processing

Complex Morlet wavelets were calculated for frequencies between 4.704 Hz and 38.416 Hz in steps of .784 Hz on single-trial data for all conditions. A Morlet constant of 7 was used throughout. Single-trial power values (evolutionary spectra) were obtained as the convolution of the wavelet family with the data and were kept for single-trial modeling and classification analyses. For visualization only, power was averaged across trials and participants per condition for illustration purposes. To address potential confounds of the mean alpha power measures with properties of the spectrum such as the overall offset, or slope of the 1/f spectral shape (e.g., Voytek et al., 2015), analyses were performed on raw alpha power as well as on residual alpha power with the 1/f shape of the spectrum removed. To remove 1/f effects, we used an approach similar to Ouyang et al. (2020), decomposing the spectrum into a 1/f portion and a specific alpha portion by fitting a second-order polynomial to the wavelet spectrum at each time point and measuring alpha as the residual power in the frequency band of interest, after removing the fitted curve through division. This results in a unitless index of signal-to-noise ratio, measuring power increases specifically in the alpha band. Both raw and 1/f corrected power was measured as the mean between 8.62—11.76 Hz, to capture the range of individuals’ peak alpha frequencies. The temporal smearing (full width at half maximum, FWHM) at the lowest frequency in this range, 8.62 Hz, was 129.5 ms and 94.7 ms at the highest frequency (11.76 Hz).

### Rescorla-Wagner Model Fitting

#### Expectancy Ratings

For each participant, we determined the RW model in which the free learning rate parameter (*α*) met the least-squares criterion for the residuals between the trial-wise standardized expectancy ratings and the associative strength V, given the empirical contingencies. The model also contained an intercept parameter to accommodate any overall offset of the ratings. Contingencies were coded as 1 for paired trials (grating plus loud noise) and 0 for unpaired trials (grating without loud noise). We also fitted a version of the Pearce-Hall model as described in the next paragraph. Models were compared regarding their data recovery and in terms of the Bayesian Information Criterion (BIC; Schwarz, 1978), which penalizes the number of free parameters. For model comparisons at the group level, BIC values were aggregated over all participants for each model. Smaller BIC values indicate better model fit. The nlinfit function implemented in MATLAB software (version 2018a) was used for the model fitting procedure, using the Levenberg–Marquardt algorithm with a limit of 10,000 iterations.

#### Alpha band oscillations

We extracted mean alpha power (8.62-11.76 Hz) from individual trials from three time-windows of interest; Post-rating, 150 to 650 ms after the start of the anticipation period, Pre-cue, 650 to 150 ms before CS onset, and Post-cue, 150 to 650 ms after CS onset. These time ranges were chosen to correspond to the three key phases of the paradigm (rating, anticipation, cue perception) while taking into account the temporal smearing at the lowest frequency included in the alpha band, i.e., 129.5 ms at 8.62 Hz. Paralleling rating data, alpha power was standardized per sensor by dividing by the maximum of the single-trial time series, separately for each time range average, and matched with the corresponding previous trial outcome. Paralleling behavioral data, we used a least-squares non-linear regression to fit the RW model (see equation 1) to the data with free parameters of learning rate (*α*) and intercept (ε) values.

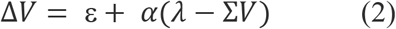

Contingencies (*λ*) were the outcomes of the previous trial, i.e., the presence or absence of a US, coded as 1 and 0, respectively. To examine the impact of separately fitting a saliency/associability parameter, we fitted a version of the Pearce-Hall model, in which a fixed learning rate was used together with the trial-by-trial adjustment of a parameter that with each trial *i* approximated the prediction error (*λ*_*i*_ − Σ*V*_*i*−1_). The Pearce-Hall (PH) model assumes that attention to the CS increases as a function of the prediction error, with greater prediction error prompting greater attention/saliency of the stimulus (Li et al., 2011). We modeled this dynamically changing attention parameter *β*_*i*_ as the product of a scaling factor *ω*, modified by the trial-wise prediction error.

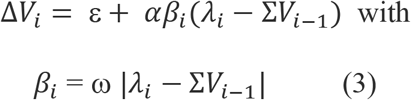

The best-fitting learning rate and saliency values were then used as parameter inputs for the same RW and PH model to produce a modeled alpha power time series for each sensor. In addition, we recorded root mean square errors and the Bayesian information criterion as metrics of the goodness of fit. When appropriate, corresponding Bayesian analyses were conducted in addition to frequentist analyses to confirm observed effects. Interpretation of Bayes factors follows the guidelines proposed by Jeffreys (1961). Predicted trial-by-trial alpha power was correlated with observed alpha power within each trial, sensor, and participant, to quantify the predictive value of the RW model on alpha power fluctuations. Bayesian information criteria for each sensor were also calculated for model comparison. This criterion determines the model fit while penalizing for increasing numbers of parameters in a given model. This makes it possible to directly compare the RW and PH model, which have different numbers of parameters.

### Binary Classification Analyses

#### Support vector machine learning

To assess the extent to which alpha oscillations contain decodable information regarding experience and expectancy, alpha power topographies were classified for each time point as a function of (1) previous trial US outcome (noise vs. no noise) and (2) US expectancy rating (upper tertile vs. lower tertile). All 129 scalp electrodes were included in the topographies making up the feature vector for each time point. The MATLAB fitclinear function which linearly classifies high-dimensional data was used. A lasso-penalty was used for classifier regularization. Classification was performed on 60 trial averages (30 for each category) with 40 trials per average, composed by randomly selecting 2 single-trial time-frequency representations from each subject, for each label. Trial averaging has been shown to heighten the robustness of classifier performance on EEG data (Bae & Luck, 2018). Here, we averaged across participants to maximize the generalizability of the patterns identified. Each topography was normalized with a z-score transformation to remove the mean, as recommended for SVM classifiers. The fitclinear function was then used to train an SVM with a 6-fold cross-validation procedure. In this process, the classifier was trained on 50 trial averages and tested on the remaining 10, for each time point separately. This was repeated until all trial averages were used as the test set. This 6-fold cross-validation was then repeated 50 times on different combinations of trial averages (using the MATLAB randperm function) to assess the variability/stability of the classification. Accuracy values were determined from the diagonal of the confusion matrix and averaged across the 6 folds and 50 repetitions. Accuracy data were then smoothed using a moving average over 20 sample points. Classifier error was then calculated by running the same training procedure on data with randomly scrambled labels, for a total of 5,000 classifications. To quantify the separation of the actual classification accuracy from the error distribution, z scores for each time point were computed by subtracting each decoding accuracy by the mean of the error distribution and dividing by the standard deviation of the error distribution.

#### Temporal generalization

All previous classifier training and testing procedures were also applied in the temporal generalization analysis. A classifier was built for each time point and then applied to each additional time point in the trial matrix using the same cross-validation procedure as described above. Pointwise classification refers to training and testing a classifier to label a topography from a given time point as one condition or another. Generalized classification refers to training a classifier on time point A and testing it on all other timepoints. The diagonal of a table or figure refers to instances when the classifier was trained and tested at the same time point. High classifier accuracy across time points indicates that the same feature patterns (classifier weights) decode the labels during those different times. The forward-modeling technique proposed by Haufe et al. (2014) was used to visualize the weight of each feature (i.e. individual sensors) in the SVM classifier at each time point. The topographical similarity between SVM weights at different time points was measured using a normalized Cartesian distance with the D statistic (Galán et al., 1997). The D statistic is a measure of the mean difference in spatial distance between corresponding pairs of two topographies, ranked by magnitude. In the present study, the sensors were ranked in order of weight value and the first 20 pairs entered the comparison between two topographies. These values were compared to a random permutation with 5,000 iterations and converted to z-scores to test for statistical significance. Smaller values indicate greater similarity. The resulting z-scores allowed us to assess topographical similarity in three graded steps: More similar than expected by chance (*z* < −2.04), as similar as expected by chance (−2.04 < *z* < 2.04), and more dissimilar than would be expected by chance (*z* < 2.04).

## Results

### Behavioral Expectancy Ratings

#### Manipulation check

Paired samples t-tests showed that participant’s standardized expectancy ratings were greater after noise (i.e., US) trials (*M* = −.4, *SD* = .313), than after no-noise trials (*M* = .411, *SD* = .31), t(19) = 5.875, p<.001, Cohen’s d = 1.314, with a corresponding Bayes factor BF_10_ = 1909.2, providing strong evidence for the notion that expectancy ratings varied as a function of recent outcome. This is consistent with the prediction of the RW model, stating that expectancy reflects recent experience, modulated by the learning rate.

#### RW model fitting

To further characterize the relation between past outcomes and updating of expectancy ratings, we fit the RW model to the expectancy time series, for each participant. At the group level, the RW model fit the data well, with the correlation between RW-predicted and observed expectancy ratings across all trials and participants being *r* = 0.47 (BIC=-362). At the level of individual participants, 12 out of 20 participants’ trial expectancy ratings were predicted at high accuracy, with Pearson coefficients between modeled and observed data greater than .14, corresponding to a p-value of 0.05. As expected, the modified Pearce-Hall model in which the stimulus saliency was learned from the contingencies fit the data better (overall *r* = 0.55; BIC=-421) than the RW model with one free learning rate parameter. Individual model RW parameter estimates for the RW and correlations between predicted and observed data are illustrated in Figure 3b, which also illustrates that the model fit and the best fitting learning rate did not systematically co-vary across participants.

**Figure 1.**
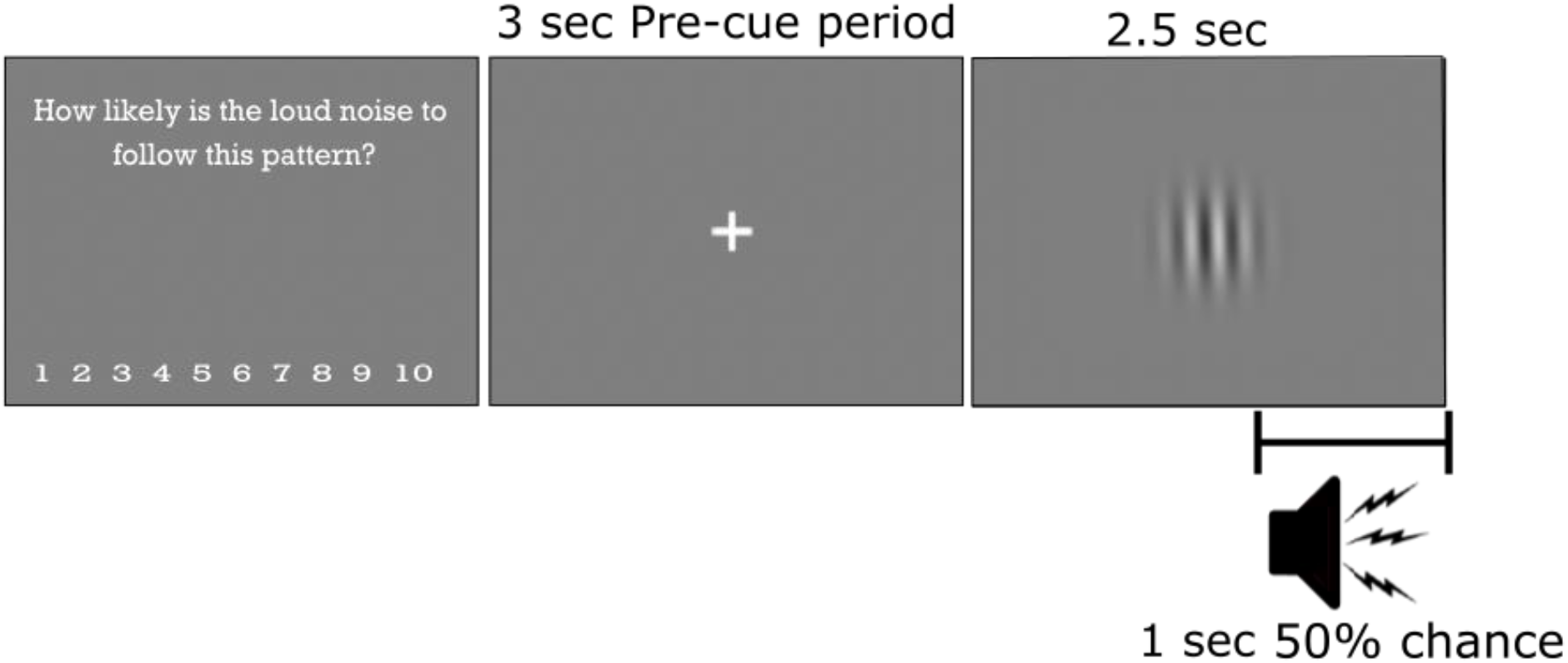
Timeline of an experimental trial. Participants rated the subjective likelihood of an upcoming US, saw a fixation cross for 3 seconds, then the orientation grating (CS) for 2.5 seconds total with a 50% chance of a loud noise (US) occurring in the last second.

**Figure 2.**
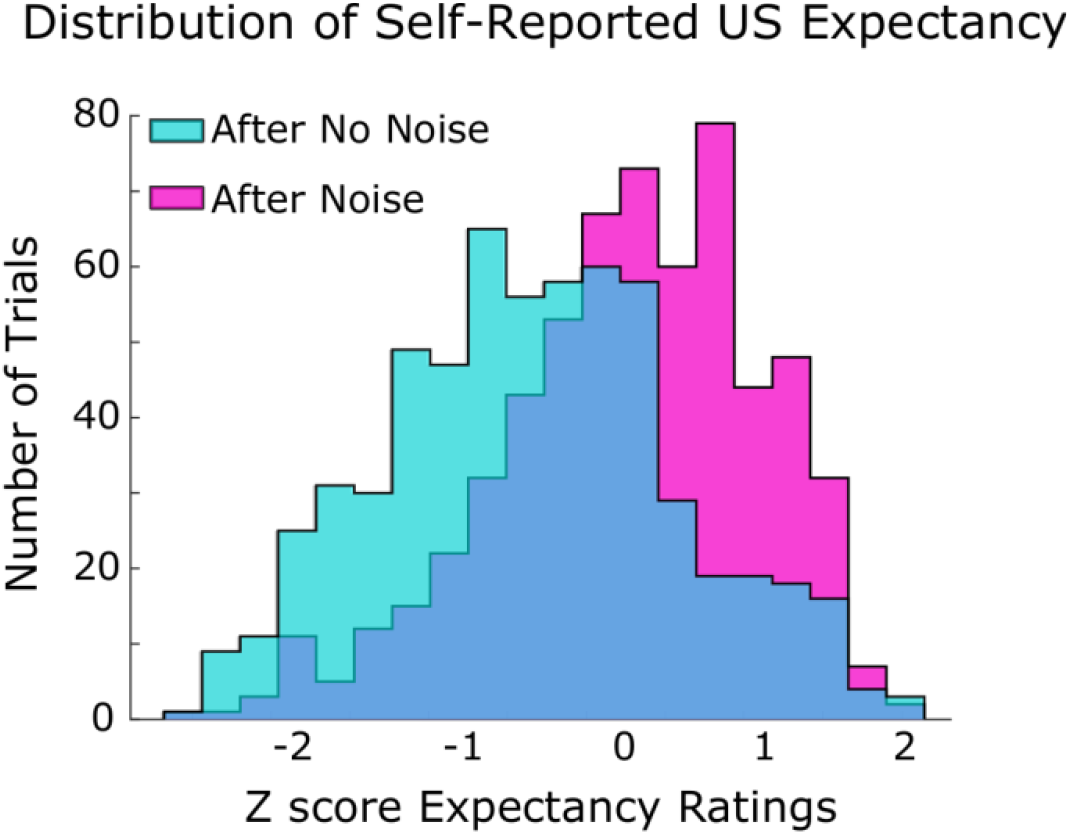
Distributions of expectancy ratings as a function of previous trial type (noise, no-noise) which occurred after a trial with a loud noise and after a trial with no noise. The expectancy of an upcoming US noise was greater after trials with a loud noise, compared to after trials in which the noise was absent.

**Figure 3.**
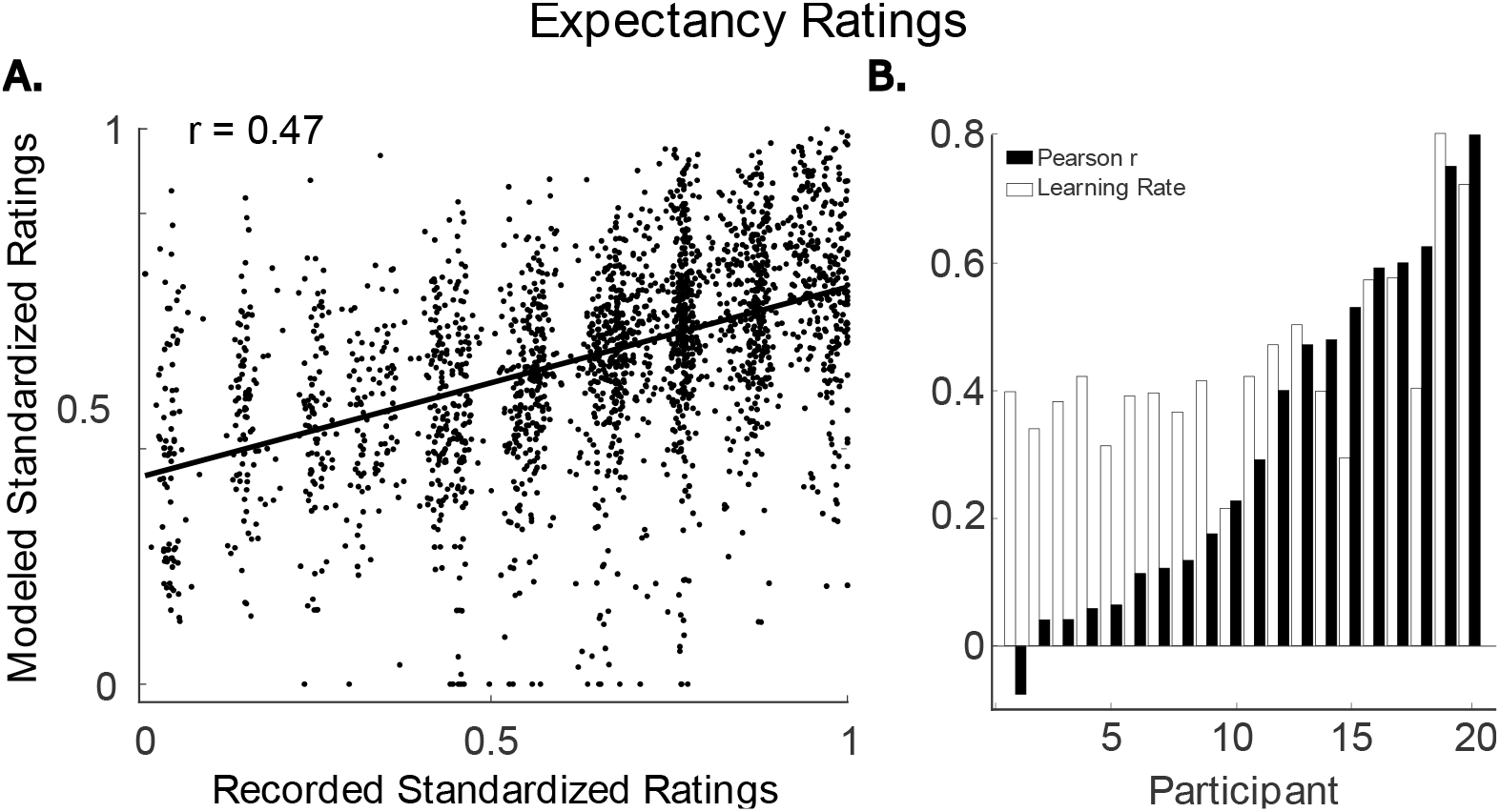
Recorded (empirical) and RW-modeled expectancy ratings for all trials and participants are plotted against each other in panel A. The grand mean Pearson correlation over all observed and modeled ratings was *r* = 0.47. Individual correlations between modeled and observed expectancy ratings for each participant are shown in panel B, along with the best-fitting learning rate for each participant.

### Neural Data

#### Descriptive

Alpha power was most pronounced during the second before the onset of the cue, followed by a decrease at the trial onset, and was characterized mostly in parieto-occipital scalp regions (see Fig.4).

**Figure 4.**
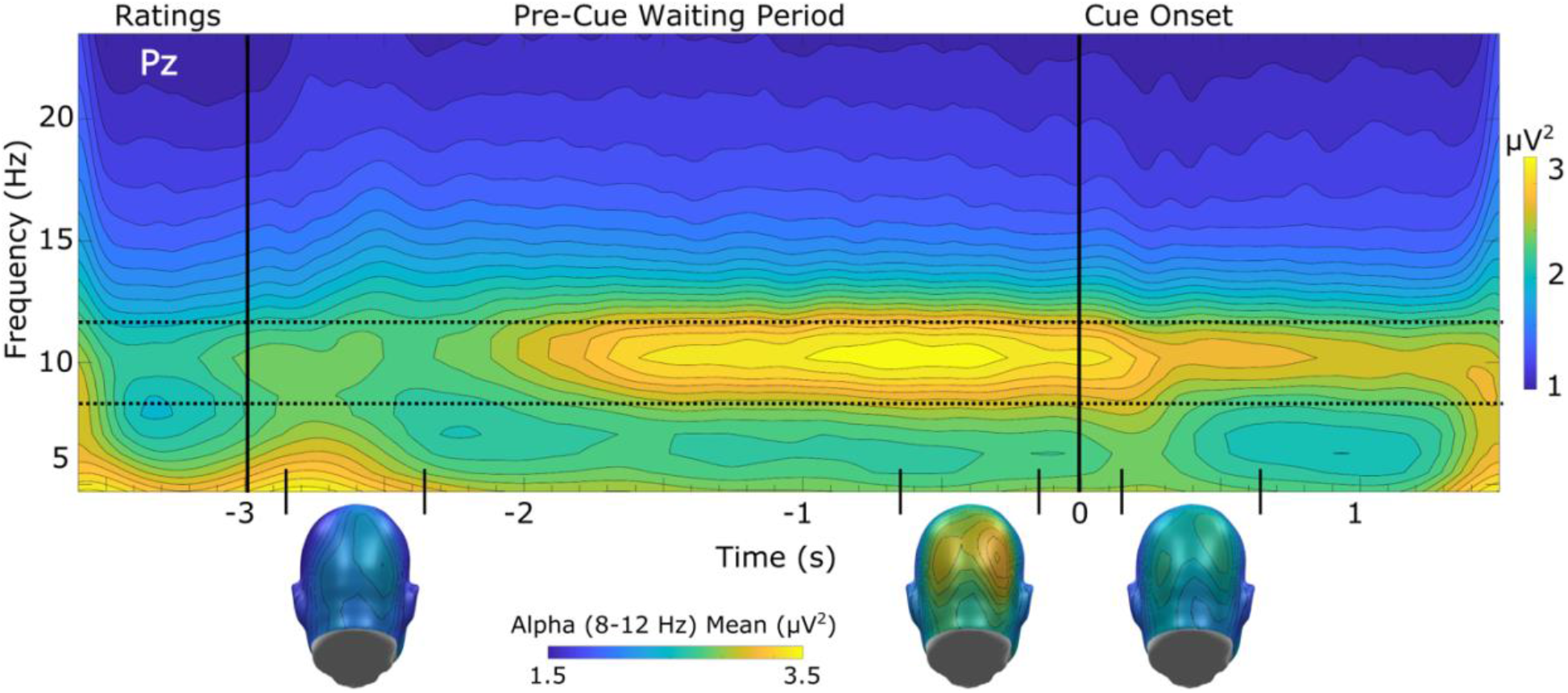
Grand mean (n= 20) time course of time-varying EEG power from a parieto-occipital sensor (EGI sensor 62; Pz) with corresponding topographies averaged over time periods of interest. Non-baseline-corrected EEG power in the alpha range (indicated by horizontal dashed lines) was lowest during and after the rating period, greatest in the pre-cue period, and showed the expected reduction after the cue onset.

#### RW model fitting

We quantified the ability of the RW model with one free parameter to capture variability in trial-by-trial alpha power during different temporal segments of the task. As expected, the RW-predicted single-trial alpha power values were most similar to observed single-trial alpha power values at parieto-occipital sensors, during the time ranges with the greatest alpha power (see Figure 4 above). Grand mean correlation maps quantifying the linear relation of predicted and observed values at each sensor of the EEG array are shown in Figure 5, for the three temporal regions of interest (post-rating, pre-cue, post-cue). Corresponding grand mean best fitting learning rate parameters for these time windows are shown in Figure 5. Across all time periods of interest, model topographies showed strong convergence: Lower learning rates co-occurred with better model-data correspondence, specifically at parieto-occipital sensors. Normalized cartesian distances between the correlation and learning rate maps showed that parameter and fit topographies were more similar than would be expected by chance; post-rating time window z = −3.4, pre-CS time window z = −4.6, post-CS time window z = −3.4. Thus, across the sensor space, better model fitting was associated with lower learning rate parameters. Correlations between model and data were overall satisfactory to very good, with a majority of individual participants, and the combined data, showing the robust fitting of the RW model (see Figure 6). The Pearson correlations between observed and predicted single-trial alpha power for each of the three time-windows across all data were as follows; post-rating time window r = .45, pre-CS time window r = .49, post-CS time window r = .43.

**Figure 5.**
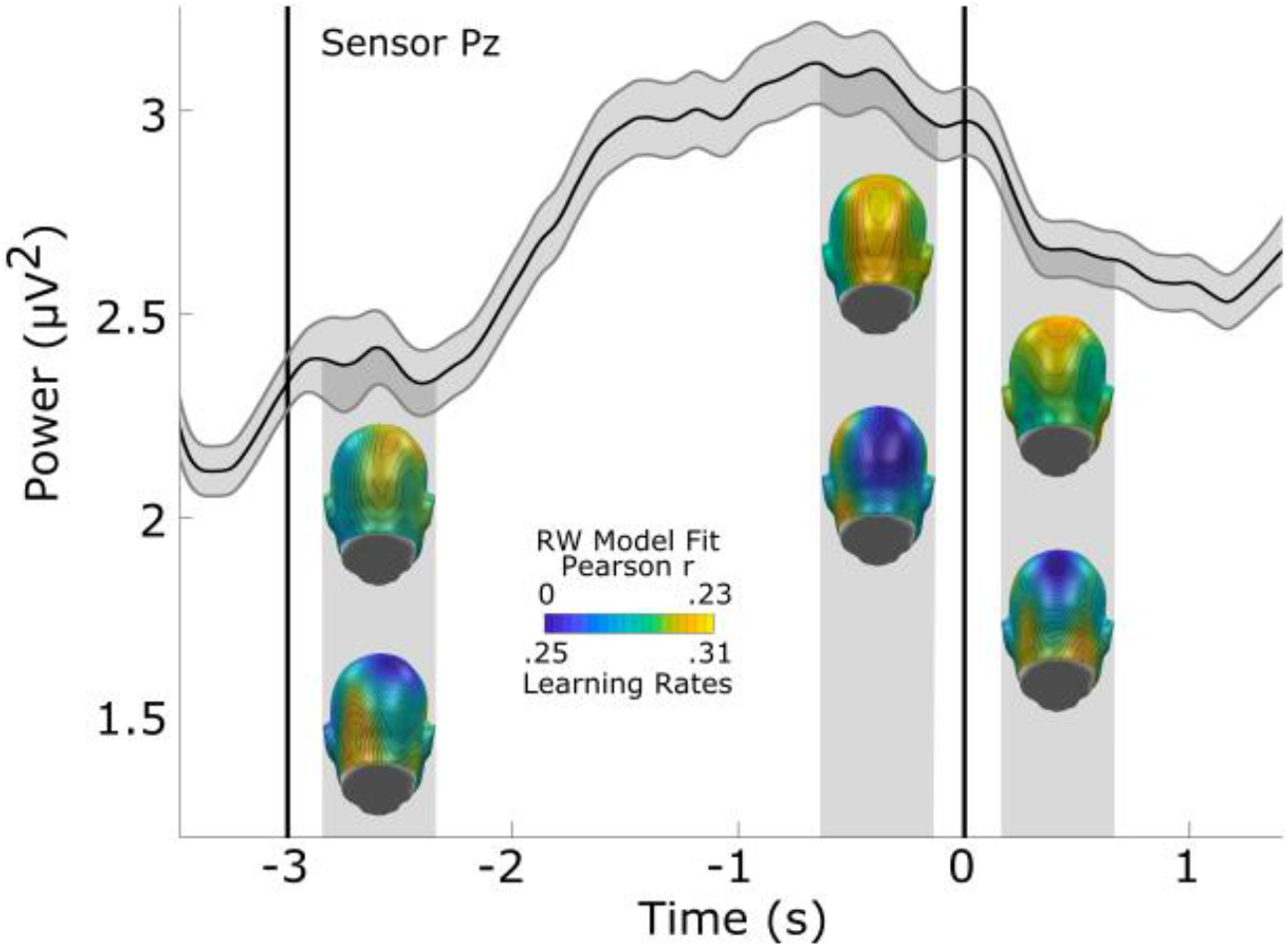
Grand mean (n=20) time-varying power in the alpha band (black line) and RW model fit (Pearson correlations between modeled and observed alpha; top topographies) and learning rates (*α*; bottom topographies) for three selected time ranges, post-rating, pre-cue, and post-cue (Pearson correlations between modeled and observed alpha). The gray shaded area displays the within-subjects standard error of time-varying alpha power for each time point, recorded at mid-parietal sensor Pz. Topographies show the grand mean Pearson correlation between model and data, as well as best fitting learning rates for each sensor.

**Figure 6.**
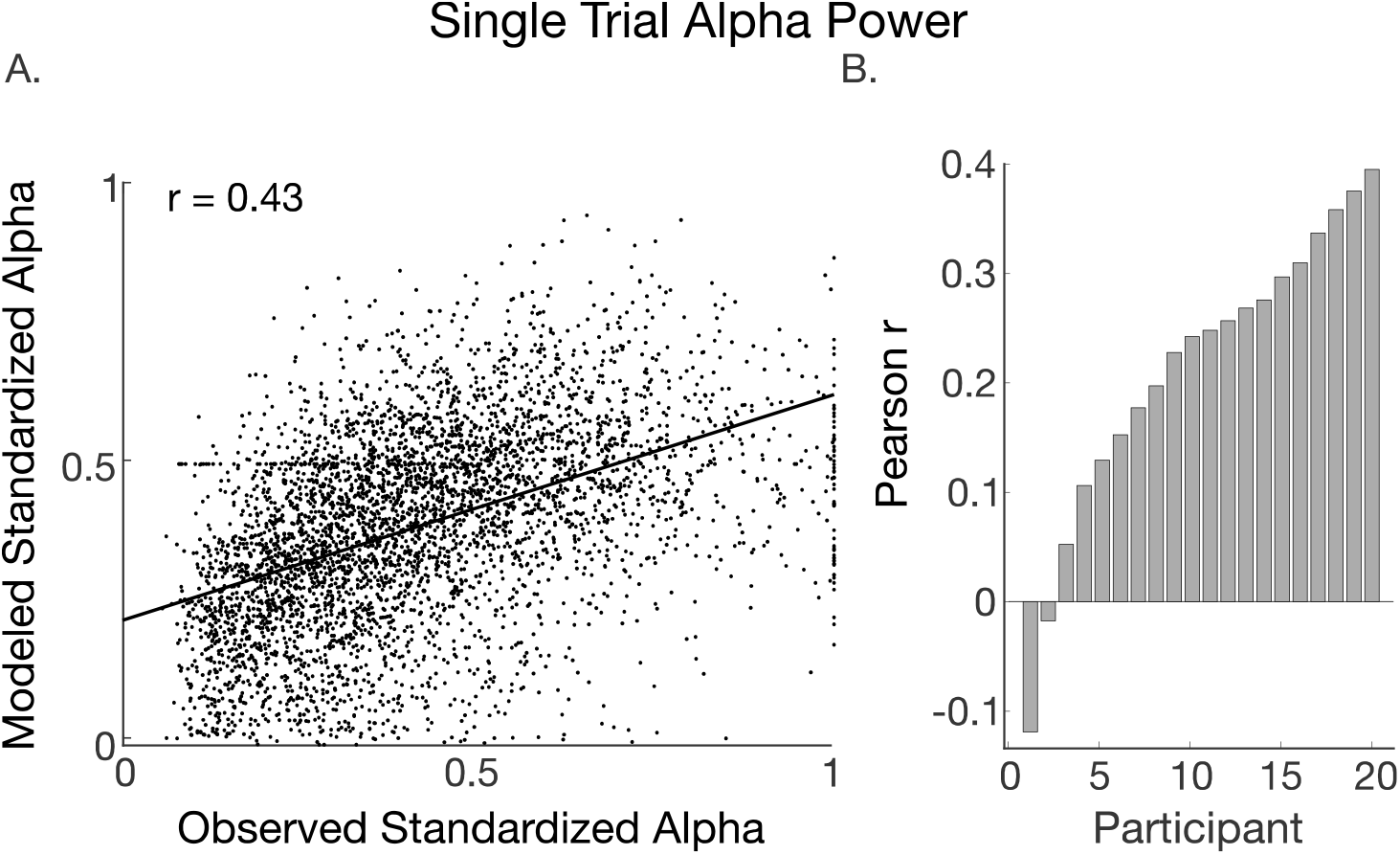
A) Scatter plot between observed alpha power and RW-predicted alpha power, across all participants and trials, averaged over the three temporal regions of interest. B) Pearson correlation coefficients measuring the fit between modeled and observed single-trial alpha data for each participant.

There were no between or within-subject relationships between behavior and alpha modeling goodness-of-fit measures. In other words, subjects who displayed high RW-fitting behavioral data did not necessarily have high RW-fitting neural data. Correlations between behavior model fit measures (Pearson r) and neural model fit measures (Pearson r) at sensor Pz for each time window were as follows; Post-Rating, *r*(19) = −.113, *p* =.64 Pre-Cue Onset, *r*(19) = −.059, *p* = .80, Post-Cue Onset, *r*(19) = −.303, *p* = .19.

Pearson correlations for the modified Pearce-Hall model were compared to RW Pearson correlations at each sensor by subtracting the PH fits from the RW fits. Differences across all sensors in each time window were between 0 and −.16. BICs comparing the models also did not systematically differ but the RW model overall showed a tendency to have lower BICs than the PH, suggesting that the alpha power data were captured well by the RW model. Thus, unlike for expectancy ratings, the trial-wise updating of saliency in the PH model did not result in a better model fit of alpha power, which was well captured by the standard RW model with one free learning rate parameter.

#### Classification Analysis: Temporal Generalization

Results from temporal generalization analyses with the SVM classifier are shown in Fig.7. Classification accuracy greater than chance as determined by permutation tests (5,000 permutations) are outlined in black. Classification accuracy along the diagonal corresponds to pointwise SVM analyses, in which two conditions are classified based on data from the same time point. Highlighted areas off of the diagonal indicate instances in which a classifier trained on topographies from a given time point accurately distinguishes the conditions during additional time ranges. In classification analyses of both US experience and US expectancy, SVMs with high accuracy during early rating and post-rating range reliably return around the onset of the CS (corresponding to time 0 in Figure 7). The classifier accuracy and temporal extent of these generalization effects were greater for classification based on expectancy than for classification based on experience (Figure 7). Topographies that resulted in the strongest classification generalization of expectancy conditions were more restricted to periods during which participants were making expectancy rating decisions with the return being strongest at visual cue onset. By contrast, in experience classification, topographies that returned at visual cue onset came from the time period immediately after the expectancy rating decision had been made and were sustained into the pre-cue period. Sustained accurate classification during the pre-cue period was observed only for decoding by expectancy but not by experience.

**Figure 7.**
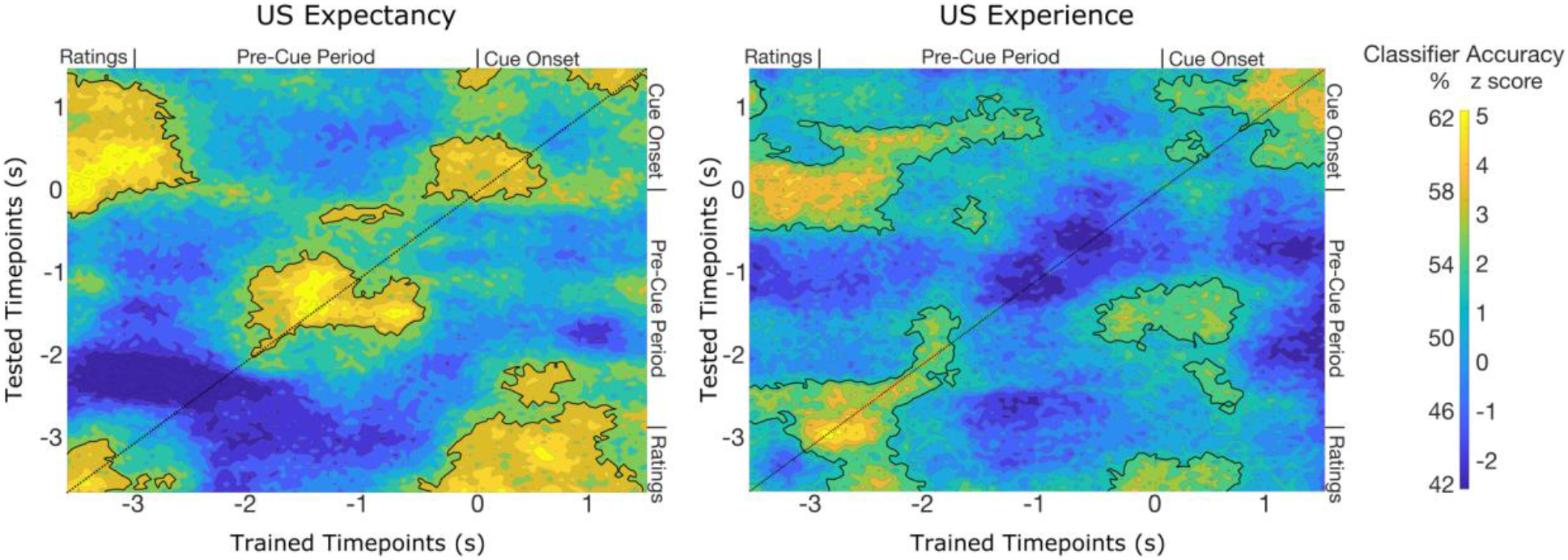
Classification accuracy represented as both percent correct and z-scores (accuracy relative to a permutation distribution) for every tested time point. Displayed along the x-axis are the time points used for SVM training, while the y-axis represents timepoints used for SVM testing. Outlined regions indicate time ranges in which classification accuracy was above chance.

Forward models of the point-wise alpha power classification were created by projecting sensor weights from the point-wise classifier back onto the scalp in Figure 8 (Haufe et al., 2014). Measures of similarity between the models are displayed at the top of Figure 8, using normalized D statistics (Galan et al., 1997).

**Figure 8.**
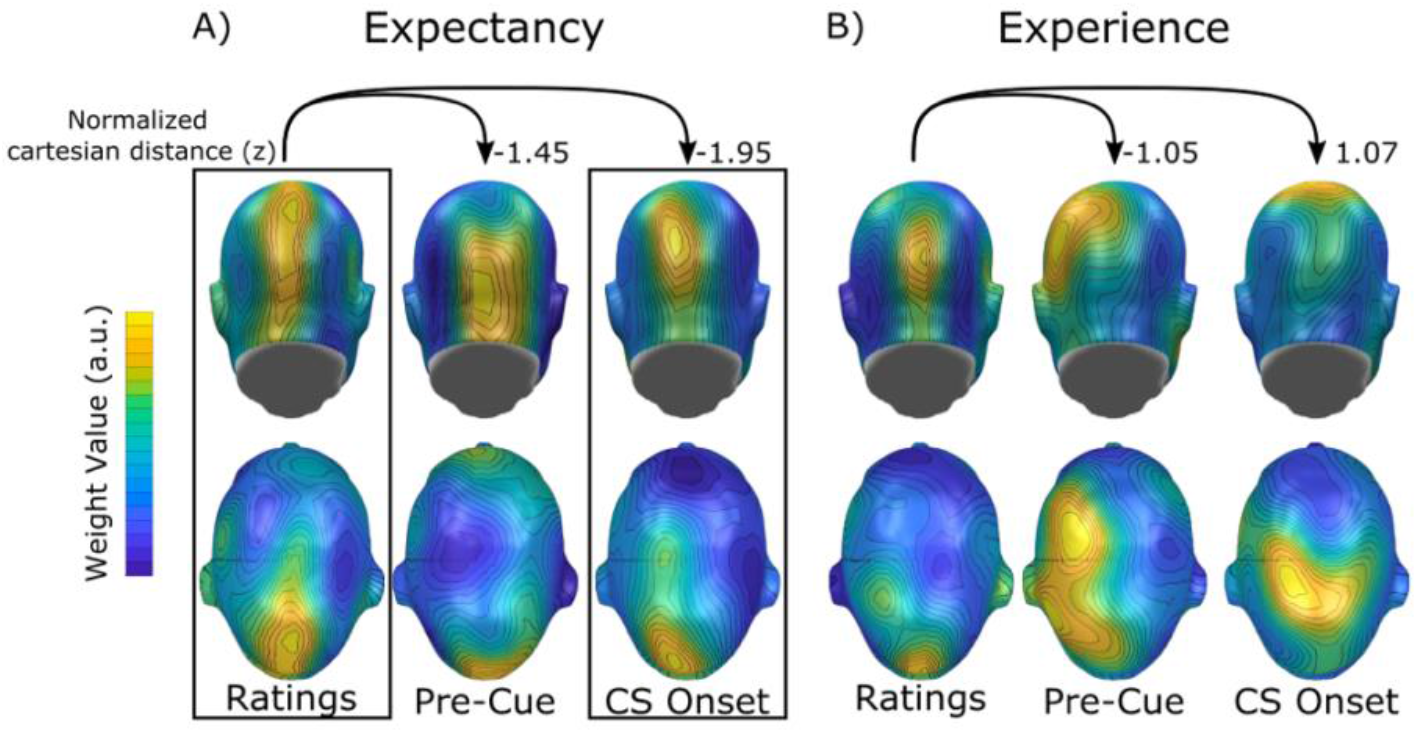
Forward-modeled SVM weight maps and their topographical similarity relative to the post-ratings time window. A) Topographical distributions of expectancy-SVM weights show maintained similarity, with the greatest weights present at parietal and occipital sensor locations throughout the trial. B) Forward-modeled weight maps for the experience-SVM (noise presence vs. absence) also show contributions from predominantly parietal and occipital sensors to the classification. Less similarity across time points is visible however along with a pronounced contribution of superior parietal sensors to the classification during cue onset.

## Discussion

The present study examined the extent to which neural oscillations in the alpha band contain information related to the trial-by-trial updating of expectations based on experience. To address this question, data were fit with a computational model (the RW model; Rescorla & Wagner, 1972) and examined with a data-driven machine learning technique. Participants reported trial-by-trial expectancy of a loud noise in a simple aversive conditioning paradigm in which a visual CS was occasionally (50 % contingency) paired with a loud noise. A waiting period followed each rating, during which participants anticipated the next conditioned cue and unconditioned stimulus.

Both subjective expectancy ratings and oscillatory power in the alpha range (8-12 Hz) were successfully modeled by the Rescorla-Wagner Learning model, with one free parameter, the learning rate. A version of the PH model, in which an additional saliency parameter was learned and updated based on each trial’s prediction error, did not further improve model performance for neural data, but did improve the fitting of behavioral (rating) data, as has been reported extensively in the literature (e.g., Diederen & Schultz, 2015). This suggests that the core mechanism of the RW model captures substantial variance in alpha power fluctuations. We also found that posterior alpha power is best characterized by RW models with lower learning rates (ranging between .3 and .5). These learning rates represent slower updating than was reported previously for primary visual cortical signals (showing a learning rate of 1; Yuan et al., 2018), and also slower updating than observed for expectancy ratings in the present study (ranging between .4 and .8), consistent with previous behavioral studies (Diederen & Schultz, 2015).

Taken together, memory updating during associative learning is tightly linked to fluctuations in the magnitude of alpha activity, and it does so at a low learning rate, consistent with integrating information over multiple trials (Gershman, 2015). Classification analyses were used to define topographical features of alpha-band oscillations that are uniquely linked to states of high versus low US expectancy, as well as to past experience (noise or no noise). Classifiers were trained pointwise and applied both within and across time points, in a temporal generalization analysis (King & Dehaene, 2014). These analyses revealed that alpha power topographies contained uniquely decodable information for both experience and expectancy.

Alpha power topographies recorded during and immediately after the rating stage of the experiment distinguished high vs. low expectancy trials. Furthermore, expectancy was decoded by the alpha power topography during the pre-cue period, starting at around 2000 ms prior to the cue, and during cue perception. Importantly, temporal generalization analyses indicated that the classifier weights from the rating stage also decoded expectancy during cue perception, for a duration of 900 ms. Forward modeling of the SVM weights indicated that the shared representation of expectancy during ratings and cue perception was based on a strong contribution of mid occipital and mid parietal sensor locations. Such topography is consistent with stimulus-induced alpha reduction, traditionally observed at posterior sites in studies of visual perception and attention (Adrian & Matthews, 1934; Klimesch, 2012).

Decoding by experience (i.e., the presence of a noise in the previous trial) was less pronounced overall but showed sustained decoding immediately after the rating period for approximately one second of time, as well as in the latter half of the cue perception period. Notably, experience was not decoded during the pre-cue interval, where expectancy-based decoding was pronounced. Temporal generalization analyses indicated that the classifier weights during and after the rating stage, a total of about 3000 ms, also decoded well during the cue onset and perception period, paralleling expectancy decoding. Comparing experience and expectancy-based decoding highlights the role of pre-cue alpha power increase in visual and parietal areas, as a specific neural substrate of expectancy formation. Such a functional role is consistent with the excellent RW fit during the same pre-cue period. Together, RW associative strength and expectancy decoding converged in their sensitivity to anticipatory processes, a well-established modulator of alpha power in other experimental paradigms such as detection tasks (Weisz et al., 2014), working memory (Bonnefond & Jensen, 2012), or cued attention tasks (Petro & Keil, 2015).

Modeled neural data from both analysis techniques were projected back onto the scalp to identify scalp locations where changes in alpha power contributed to the modeling and classification analyses. The resulting weight maps showed strong support for the notion that the spatial configuration, as well as the time-varying power in posterior alpha-band activity, both contain unique information that is tightly linked to the experience-based updating of expectancy. As an overview, Figure 9 shows the model fit and SVM weight topographies for each analysis method, along with non-parametric tests of mean spatial distance between the topographies. In these comparisons, none of the topographies were more similar than would be expected by chance, and only the difference between the RW model fit and the US experience classifier topographies was more different than would be expected by chance. This latter fact is again consistent with the notion that expectancy decoding and RW modeling only partially overlapped in their spatial configuration. The topographical comparisons of weight maps support the notion that oscillatory activity in the alpha band is not monolithic but may comprise multiple parallel processes (Ben-Simon et al., 2008), distinguished by their source configuration and additional features such as phase or waveform shape (Klimesch et al., 2000; Mathewson et al., 2011). In the present study, each analytical approach portrays aspects of alpha that contribute to different facets of associative learning on a trial-by-trial basis.

**Figure 9.**
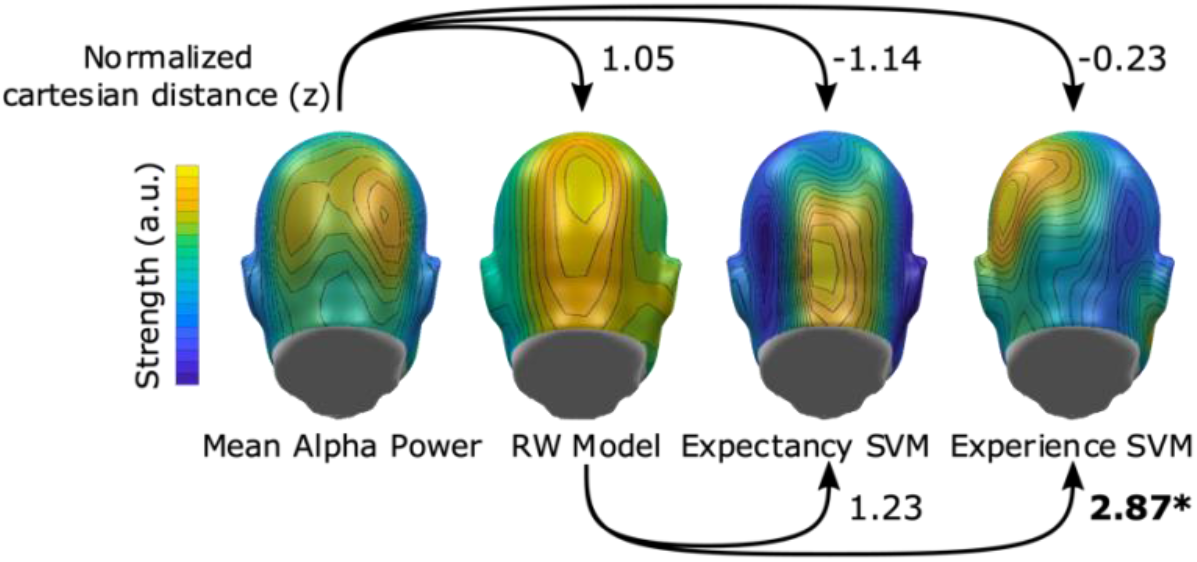
Measures of topography similarity between data and different analyses during the pre-cue time segment. Topographical differences in the tails of the permutation distribution (Zs exceeding ± 2.04) are considered more similar (negative values) or more different (positive values) than would be expected by chance.

Temporal generalization analyses of both expectancy and experience showed that the decodable information present during and after the rating also correctly classified the labels during and after the onset of the conditioned cue. This finding, together with the occipital topography of the weights, would be in line with observations that representations of contingencies between a visual cue and an outcome have a strong visuocortical component (Moratti & Keil, 2009; Shuler & Bear, 2006). It is also consistent with the notion that cognitive processes of anticipation and perception overlap neurophysiologically (Klimesch, Sauseng, & Hanslmayr, 2007), perhaps resulting in a partial replay of the rating-related neurophysiological cascade upon viewing the visual cue (Ekman, Kok, & de Lange, 2017). Other hypotheses regarding the role of alpha in perception and memory formation include (i) that alpha power modulations first act as a perceptual focusing mechanism and, through attention-like mechanisms, affects subsequent memory retrieval (Wang, Megla, & Woodman, 2020), and that (ii) multiple mechanisms use the alpha frequency band to encode and maintain current stimulus representations for prospective use (de Vries et al., 2017). Broadly in line with these notions, the present findings demonstrate that just one model, or proposed function, is incapable of capturing the role of alpha-band oscillations. Multiple spatial and temporal configurations, along with magnitude changes had unique correlates within the association formation cycle. Studies in awake non-human primates show that the neural generators of alpha rhythms may show complex and even opposite alpha power and frequency changes across neural regions (Bollimunta et al., 2008)—further support for emerging neurophysiological models in which alpha changes are seen as reflecting changes in the excitability or gain sensitivity in specific tissues relevant to a given task (Benwell et al., 2019). Such a view is inconsistent with mapping band power changes to one cognitive process, and instead emphasizes the relevance of the specific task, along with the temporal and spatial configuration of alpha power changes, afforded by modeling and multivariate decoding analyses as employed here.

There was no benefit of using the flexible PH model over the more static standard RW model when fitting trial-wise alpha dynamics in the present study. Previous work has identified specific brain regions, which are best modeled by PH versus RW models, and which are reflective of different variables mapped onto concepts such as salience, attention, and prediction error (Roesch et al., 2012). At the broad level of scalp EEG, such specificity is unavailable, and thus, the lack of support for a PH mechanism may be a reflection of using scalp EEG to capture broad post-synaptic neural communication at the network level (Nunez & Srinivasan, 2006). However, a growing body of research shows that EEG measures are sensitive to prediction error during certain types of learning (Burnside, Fischer, & Ullsperger, 2019) and that arousal system responses are overall more predictive of surprise (de Gee et al., 2020), suggesting that the present findings cannot be explained by the limitations of EEG alone.

### Conclusion

The present study demonstrates that time-varying changes in EEG alpha power contain information that is uniquely linked to trial-by-trial updating of expectancies during associative learning. Despite pronounced inter-individual variability, the modeling and decoding approaches used here converged in showing that expectancy ratings were strongly linked to specific oscillatory states, reflected in power fluctuations and the topographical configuration of the alpha signal. As such, the present analyses provide more information than trial averages, and univariate analyses, paving the way to using behavioral and neural measures as meaningful indices of learning (Haines et al, 2020). Such future work may use more advanced modeling techniques for integrating behavioral and neural data directly to further define the multiple, unique roles of alpha oscillations in the formation of associative memories and the subjective experience of anticipation.

